# Linear β-1,2-glucans trigger immune hallmarks and disease resistance in plants

**DOI:** 10.1101/2024.05.30.596602

**Authors:** María Fuertes-Rabanal, Asier Largo-Gosens, Alicia Fischer, Kristina S. Munzert, Cristian Carrasco-López, Andrea Sánchez-Vallet, Timo Engelsdorf, Hugo Mélida

## Abstract

Immune responses in plants are triggered by molecular patterns or elicitors, recognized by plant pattern recognition receptors (PRRs). Such molecular patterns arise from host-pathogen interactions and the response cascade activated after their perception is known as pattern-triggered immunity (PTI). Glucans have emerged as key players in PTI, but certain glucans’ ability to stimulate defensive responses in plants remains understudied. This work focused on identifying novel glucan oligosaccharides acting as molecular patterns. The ability of various microorganism-derived glucans to prompt PTI responses was tested, revealing that specific microbial-derived glucans, such as short linear β-1,2-glucans, trigger this response in plants by increasing reactive oxygen species (ROS) production, MAP kinase phosphorylation, and differential expression of defence-related genes in *Arabidopsis thaliana*. Pretreatments with β-1,2-glucan trisaccharide (B2G3) improved Arabidopsis defence against bacterial and fungal infections in a hypersusceptible genotype. The knowledge generated was then transferred to the monocotyledonous model species maize and wheat, confirming that these plants also respond to β-1,2-glucans, with increased ROS production and improved protection against fungal infections following B2G3 pretreatments. In summary, as with other β-glucans, plants perceive β-1,2-glucans as warning signals and stimulate defence responses against phytopathogens.

**Highlights:** We describe a new group of glycans present in the extracellular matrices of some plant-interacting microorganisms that are sensed by host surveillance systems and enhance the plant’s natural resistance to disease.

## Introduction

The plant immune system is a complex network that operates in all plant cells, unlike mammals that have specialized immune cells (DeFalco and Zipfel, 2021). Plants have evolved a multifaceted immune system that involves different layers of stimulus perception and defence mechanisms (Yuan et al., 2021). One of the essential layers of defence in the plant-pathogen interaction is the plant cell wall, which acts as a physical barrier and can be remodelled as a defensive response (Bacete et al., 2018; Molina et al., 2021). Another critical layer of plant immunity is based on the perception of conserved associated molecules by plant pattern-recognition receptors (PRRs). These molecular patterns can be microbe-associated molecular patterns (MAMPs), which are conserved molecules present in microbes, or damage-associated molecular patterns (DAMPs), which are plant degradation products derived from the infection (Cheung et al., 2020). PRRs are primarily located on the surface of the plasma membrane and upon recognition of DAMPs/MAMPs, trigger a cascade of responses known as pattern-triggered immunity (PTI), which may be sufficient to repel pathogen attack and tissue infection. Upon the perception of DAMPs or MAMPs, intracellular increases in Ca^2+^ and H_2_O_2_ bursts occur, which activate downstream signalling networks regulated by plant mitogen-activated protein kinases (MAPKs) and calcium-dependent protein kinases (CDPKs) (Bigeard et al., 2015). These MAPK- and CDPK-dependent signaling networks play specific roles in controlling the activities and synthesis of a diverse array of transcription factors, enzymes, hormones, peptides, and antimicrobial compounds, all of which contribute to plant resistance to pathogens (Tena et al., 2011). Furthermore, after recovering from biotic stresses plants can acquire higher levels of resistance due to the long-lasting defence responses, known as immune priming, which can persist for hours or days after pathogen detection, enabling plants to defend themselves and neighbouring plants against pathogens (Cooper and Ton, 2022).

The field of plant immunity has historically focused on peptide-triggered immunity. However, recent advances have led to the discovery of numerous glycans derived from plants, from extracellular polysaccharides of bacteria or from pathogen cell wall that induce plant immunity (Willmann et al., 2011; Liu et al., 2012; Benedetti et al., 2015; Jiang et al., 2016; Souza et al., 2017; Claverie et al., 2018; Mélida et al., 2018, 2020; Rebaque et al., 2021; Zarattini et al. 2021; Yuan et al., 2022; Fernández-Calvo et al., 2024. Among the whole spectrum of cell wall polymers present across various organisms, glucans stand out as one of the most vital types. These polymers are characterized by a glucose backbone, with variations in branching patterns, with linear glucans being the most prevalent (Synytsya and Novak, 2014). The specific glucose carbons engaged in the polymerization process play a pivotal role in shaping the biochemical properties of glucans. This specificity allows for the differentiation of a diverse array of structures such as β-1,4-glucans responsible for cellulose formation, β-1,6-glucans characteristic of fungi and some algae and β-1,3-glucans or β-1,3/1,4-glucans present in plants, algae and fungi (Burton and Fincher, 2009; Schulze et al., 2016; Souza et al., 2017; Mélida et al., 2018; Ruiz-Herrera and Ortiz-Castellanos, 2019). The high glucans diversity of both plants and their microbial pathogens, leads to a large variety of glucoligands with potential affinity for plant receptors. Nevertheless, recent evidence suggests that filamentous microbes, such as fungi or oomycetes, have evolved towards evasion of glycan-triggered immunity, highlighting the importance of understanding the intricacies of the plant immune system (Irieda et al., 2019; Chandrasekar et al., 2022). Besides, there are still different glucans, such as those linked by α-1,3 or β-1,2 bonds contained in the extracellular matrixes of some bacteria and fungi, which far from having the capacity to trigger PTI have been proposed to act as plant or mammal immunosuppressants, a phenomenon barely described in this scientific field (Rigano et al., 2007; Rappleye and Goldman, 2008; Fujikawa et al., 2012).

This revelation underscores the complexity of plant-pathogen interactions and highlights the importance of understanding glycan-triggered immunity modulation in plant defence mechanisms. Moreover, the emergence of immune priming using glycans from various sources has demonstrated efficacy in enhancing protection across different pathosystems, showcasing the potential of glycan-based technologies in plant defence (Klarzynski et al., 2000; Aziz et al., 2007; Gravino et al., 2015; Claverie et al., 2018; Rebaque et al., 2023). This concept has spurred the development of crop protection technologies focused on enhancing the natural resistance mechanisms of the plants to combat plant pathogens, promoting sustainable agricultural practices and reducing environmental impact (Conrath et al., 2015). Notably, the utilization of β-1,3-glucans and chitin as plant immunity inducers has been successful, leading to the development and commercialization of products already being applied to the fields.

Therefore, the aim of this work was to investigate the ability of plant cell defence systems to perceive glucans that have not been studied so far, such as β-1,2-glucans, and their ability to protect plants by immune priming. To this end, we investigated the ability of *Arabidopsis thaliana* (Arabidopsis) to induce immunity and to protect against microbial infections upon treatment with β-1,2 glucans. Finally, with the purpose of testing whether β-1,2-glucans could be used to protect crops, we analysed their efficacy in inducing H_2_O_2_ production and in pathogen-protection assays in monocots.

## Material and Methods

### Plant material and growth conditions

The Arabidopsis ecotype used in this study was Columbia-0 (Col-0). Seedlings used for reactive Oxygen Species (ROS), MAPK phosphorylation and gene expression assays were grown in 24-well plates (approximately 10 seedlings per well) for 8-12 days under long day conditions (16 hours light/8 hours dark) at 26°C/24°C in liquid half-concentrated Murashige and Skoog (MS) medium. For bacteria and *Botrytis cinerea* protection analyses, Arabidopsis plants were grown in a soil-vermiculite (5:1) mixture in a growth chamber under 12 hours of light/12 hours of darkness at 23°C/21°C with a humidity of 70% for 4 weeks. For *Colletotrichum higginsianum*, Col-0 and the hypersensitive mutant *pgm* were grown in a soil-clay (4:1; Fruhstorfer soil type P, Hawita, Germany:Liadrain clay, Liapor, Germany) mixture in CLF (Wertingen, Germany) GroBank growth chambers with 12 hours of light/12 hours of darkness at 22°C/20°C for 2 weeks and subsequently transferred into long day conditions (16 hours of light/8 hours of dark; 22°C/20°C) for another 2 weeks. The photon flux density was kept at 110 µmol m^-2^ s^-1^ and the plants were fertilized 7 days prior to infection with Wuxal Super fertilizer (Aglukon, Germany).

Maize (*Zea mays* cv Mikado) seeds were soaked with tap water for 4 hours and germinated in germination trays for two days at 28°C. Seedlings were transferred to square pots with P-type soil (Fruhstorfer, Hawita, Germany) and grown under 14 hours light/10 hours darkness at 28°C/20°C and 400 µmol m^-2^ s^-1^ photon flux density for 14 days in Percival (Perry, USA) PGC-105 growth chambers. Maize plants were fertilized every two days with Hoagland solution.

Wheat plants (*Triticum aestivum* cv Titlis) were grown in square pots (11×11×12 cm) with soil under long day conditions at 18°C (light) and 15°C (dark) with a humidity of 60%. Seven-day-old plants were fertilized with 2 litres of fertilizer solution per 15 pots (5 ml/L, COMPO Universal Liquid Fertilizer, Germany).

### Plant pathogenic bacteria and fungi growth conditions

*Pseudomonas syringae* pv. *tomato* (DC3000) was kindly provided by Dr. Rubén Alcazar from *Universidad de Barcelona* (Spain). DC3000 was grown in King’s Basal medium (KB; 2% (w/v) protease peptone, 0.15% (w/v) MgSO_4_, 0.15% (w/v) KH_2_PO_4_, and 1.5% (v/v) glycerol) with 25 µg/mL of rifampicin and incubated at 28°C for three days before infection assay.

*Botrytis cinerea* was kindly provided by Víctor Flors from *Universitat Jaume I* (Spain). *B. cinerea* was grown in potato dextrose agar (PDA) medium at 25°C in dark until the formation of the conidiophores. Spores were collected by washing PDA plates with autoclaved tap water, filtered through two layers of sterile gauze and counted using a Thoma chamber. Spores were diluted to 7×10^6^ spores/mL and stored as glycerol stocks. Later, for infection experiments these spores were centrifugated at 3,000 *g* for 5 minutes and resuspended in Potato-Dextrose Broth (PDB) medium. Then, spores were counted in a Thoma chamber and were diluted with more PDB to achieve a concentration of 1×10^6^ spores/mL.

*Colletotrichum higginsianum* MAFF305635 was grown in oatmeal agar medium (OMA, 5% (w/v) grinded organic oatmeal and 1.2% (w/v) agar) for 7 days at 22°C under long day conditions to promote the conidia development. Conidia were collected by washing plates with sterile distilled water and spores were counted in a Neubauer chamber. Spores were diluted to 2×10^6^ spores/mL for infection experiments with distilled water.

*Colletotrichum graminicola* (Ces.) Wils., CgM2 isolate was grown on OMA medium (5% (w/v) oat bran and 1.2% (w/v) agar) for 14 days at 26°C under long day conditions to allow conidia formation. Spores were collected by washing plates with sterile distilled water and spores were counted in a Neubauer chamber and diluted to 2×10^4^ spores/mL.

*Zymoseptoria tritici* (Swiss strain ST99CH_3D7) spores were incubated in 50 mL of Yeast Peptone Dextrose (YPD; 1% (w/v) yeast extract, 2% (w/v) peptone and 1% (w/v) dextrose, supplemented with 50 µg/mL Kanamycin) for 6 days at 18°C and rotary shaking at 120 rpm. Spores were filtered through sterile gauze, pelleted by centrifugation at 3,273 *g* for 15 minutes and resuspended into 15-25 mL of sterile water. Spore concentration was calculated using a KOVA Glasstic counting chamber (Hycor Biomedical, Inc., California) and adjusted to 1×10^7^ spores/mL.

### Carbohydrates used in the experiments

The oligosaccharides α-1,3-glucan trisaccharide -A3G3-(α-1,3-(glucose)_3_), hexaacetyl-chitohexaose Chi6 (β-1,4-(N-acetylglucosamine)_6_), MLG43 (β-1,3, β-1,4-(glucose)_3_), β-1,2-glucan trisaccharide B2G3 (β-1,2-(glucose)_3_),), β-1,2-glucan hexasaccharide B2G6 (β-1,2-(glucose)_6_), β-1,2-glucan polysaccharide (β-1,2-(glucose)_n_) were acquired from Megazyme (Wicklow, Ireland).

### Reactive oxygen species production analysis

Eight-day-old Arabidopsis seedlings grown on liquid ½ MS medium were used for this experiment. One seedling per well was placed in a white 96-well plate, then they incubated overnight with 150 μL of distilled water at room temperature. The day after, distilled water was replaced by 100 μL of 10 mM luminol (#120–04891; FUJIFILM Wako Pure Chemical Corporation) and 1 mg/mL of horseradish peroxidase (#P6782; Sigma) and incubated for two hours. Subsequently, seedlings were treated with water (mock) and the oligosaccharides described above in order to quantify the production of H_2_O_2_ using the luminol assay and a Varioskan Lux microplate reader (Thermo Scientific). For dose-response assays, different concentrations of B2G3 (from 0.01 µM to 1mM) were used.

In case of maize, 4-week-old plants at the V3 stage (with 5 or 6 developed leaves) were used, following a protocol similar to that used in Arabidopsis.

Eight disks (12.6 mm^2^) from second leaves of 2-week old wheat plants were incubated with 100 µL of 15 µg/mL Peroxidase from horseradish (Sigma-Aldrich, P6782) and 150 nM luminol L-012 for 16 h in the dark at 15 _. Thereafter, 50 µL of 300 or 1200 µM B2G3, 3 µM flg22 or distilled water (mock) were added, and luminescence was measured using Varioskan Lux microplate reader.

In all the cases, from the data obtained in the ROS accumulation analysis, the total areas under the curves were integrated using the SkanIt software (Thermo Scientific).

### Immunoblot analysis of MAPK activation

Twelve-day-old Arabidopsis seedlings grown on liquid ½ MS medium in 24-well plates were treated with water (mock), MLG43, B2G3 and B2G6 for 0, 5, 15, 30 and 60 minutes. Subsequently, seedlings were frozen in liquid nitrogen and homogenized by pestles. Protein extraction was performed with 50 µL of extraction buffer (25 mM Tris-HCl pH 7.8, 75 mM NaCl, 15 mM Egtazic acid (EGTA), 10 mM magnesium chloride, 15 mM sodium β-glycerophosphate pentahydrate, 15 mM bis(4-nitrophenyl) phosphate, 1 mM 1,4-dithiothreitol, 1 mM sodium fluoride, 0.5 mM sodium orthovanadate, 0.5 mM phenylmethylsulfonyl fluoride, 0.1% (v/v) Tween 20 and protease inhibitor cocktail (#P9599; Sigma). Total proteins were quantified by Bradford reagent (Bio-Rad). Equal amounts of proteins were separated by SDS-PAGE and then transferred onto nitrocellulose membranes which were blocked with Protein-Free Blocking Buffer (PFBS; Thermo Scientific) for 2 hours at room temperature in agitation. The membranes were incubated overnight in agitation at 4°C with Phospho-p44/42 MAPK (Erk1/2) (Thr202/Tyr204) (#4370; Cell Signaling Technology) in a dilution 1:1000 with PFBS. Afterwards, membranes were washed three times with Tris-Buffered Saline that contains also 0.1% (v/v) Tween 20 and then incubated with horseradish-peroxidase-goat anti-rabbit polyclonal secondary antibody (#10035943; Fisher) diluted 1:250 with PFBS. Finally, the membranes were developed using ECL Western Blotting Substrate (Thermo Scientific). Additionally, the membranes were stained with Ponceau S solution to evaluate equal loading (Thermo Scientific).

### Gene expression analyses

Twelve-day-old Arabidopsis seedlings grown on liquid ½ MS medium in 24-well plates were treated with water (mock), Chi6, MLG43, B2G3 and B2G6 for 0 and 30 minutes. Total RNA was extracted using the RNeasy Plant Mini Kit (Qiagen) following the protocol of the manufacturer. RT-PCR was carried out using the High-Capacity RNA-to-cDNA Kit (Applied biosystems). Reactions for the quantitative PCR were made using 2X PowerUp SYBR Green Master Mix (Applied Biosystem) using the cycling mode described in the Quick Reference of the SYBR Green Mix. The quantitative PCR was performed in Step One Plus real-time PCR system (Applied Biosystem). We used specific primers (Supplementary Table S1) for the amplification of the immune-related genes *CYP81F2* (*At5g57220*) and *WRKY53* (*At4g23810*). The expression of each gene was normalized to *UBC21* (*At5g25760*) levels.

### Arabidopsis protection assays

For Arabidopsis protection experiments against DC3000, plants were grown in a soil vermiculite (5:1) mixture in 1 L glass culture vessels jars (PhytoTech Labs) (3 plants/vessel) formerly sterilized by autoclaving and in the growth conditions described previously. Four-week-old Arabidopsis plants were sprayed with 1 mL per plant of water (mock) or B2G3 solution (500 µM), both treatments containing 0.05% (v/v) of Tween 24 (Croda) as adjuvant. The infection with DC3000 was carried out 48 hours after pretreatments. Plants were sprayed with 2 mL of the bacterium suspension (with an optical density of 0.1 at 600 nm) that contained Silwet L-77 (PhytoTech Labs) as a surfactant. Plants were maintained at high humidity for 3 hours by using a cover sprayed with water. In order to calculate the Colony Forming Units (CFUs), two leaf discs were collected from each plant at 0 (3 hours) and 3 days post-infection (dpi) and were homogenized with a pestle for recovering the bacteria that had penetrated the tissue. CFU-per-foliar-area were determined after plating serial 1:10 dilutions of each recovered leaf-extract on KB plates with rifampicin (25 µg/mL) incubated at 28°C. Pictures of the DC3000 infection symptoms were also obtained throughout the experiment.

For *B. cinerea* inoculations, Arabidopsis plants were grown in pots (one plant/pot) for 4 weeks in the same conditions as described before. Each plant was sprayed with 0.5 mL water (mock) or B2G3 (500 µM), both supplemented with 0.05% (v/v) Tween 24 (Croda) and infection with *B.cinerea* was performed 48 hours after the treatment. *B. cinerea* spores were obtained as explained previously. Fungal infection was performed by placing four 5-µL drops, containing approximately 5,000 spores, per leaf in at least three leaves per Arabidopsis plant. Plants were maintained at high humidity by using a cover sprayed with water. After 24 hours of *B. cinerea* inoculation, nine leaves from different plants were cut, weighed, frozen under liquid nitrogen and stored at -80 °C for *B. cinerea* genomic DNA quantification. *B. cinerea* quantitation on infected leaves was performed by quantitative PCR by extracting fungal genomic DNA using NZY Plant/Fungal gDNA Isolation Kit (Nzytech, Lisboa, Portugal). Quantitative PCRs were performed as described above in QuantStudio1 equipment (Thermo Fisher Scientific), using the *B. cinerea*-specific primers (Supplementary Table S1) for amplification of *β-Tubulin* gene (Bc*TUB*). In QuantStudio 1 the fungal genomic DNA quantification was quantified in relation to the fresh weight (mg) of infected Arabidopsis leaves. After another 24 hours, the remaining leaves were photographed and the lesion area was analysed using ImageJ (Schneider et al., 2012).

The ability of B2G3 to protect against *C. higginsianum* was evaluated in Arabidopsis Col-0 plants and in the starch-deficient hypersusceptible plastidic *phosphoglucomutase* (*pgm*) mutant (Engelsdorf et al., 2013). Plants were grown in pots with a soil:clay (4:1) mixture for 4 weeks according to the growth conditions mentioned before. Plants were sprayed with 0.5 mL per plant of distilled water (mock) or 1 mM B2G3 both supplemented with 0.05% (v/v) Tween 24 (Croda) and were maintained in high humidity by using a cover sprayed with water. *C. higginsianum* was inoculated 48 hours after the treatments. *C. higginsianum* conidia solution containing 2×10^6^ conidia/mL and 1 mM B2G3 in distilled water was used to perform this infection assay, while mock treatments contained 1 mM B2G3 in distilled water. To obtain a homogeneous *C. higginsianum* inoculation, 1 mL of conidia suspension were evenly sprayed on the leaf surface and plants were maintained at high humidity by using a cover that was sprayed with water to support the fungal infection. Leaf discs were collected at 3.5 dpi to quantify *C. higginsianum* genomic DNA, frozen in liquid nitrogen and stored at -80 °C. Leaf discs were homogenized using metal beads in a Retsch mixer mill with a frequency of 20 Hz for 1 minute. The extraction of fungal genomic DNA was performed using NucleoSpin® Plant II Kit (Macherey-Nagel, Dueren, Germany) following the instructions of the manufacturer. *C. higginsianum* genomic DNA quantitation was performed by quantitative PCR using Biozym Blue S’Green qPCR Kit (Biozym Scientific, Germany) in a CFX RT-PCR detection system (Bio-Rad, USA) and specific primers for Ch*TrpC* gene (Supplementary Table S1). The relative quantity of fungal DNA was normalized to leaf area (cm^2^).

### Monocot protection assays

Maize protection assays were performed against the pathogen *C. graminicola.* Maize plants were grown for 14 days, and fully expanded fourth leaves were treated with 0.5 mL per plant of water (mock) and 1 mM of MLG43 or B2G3. All solutions were supplemented with 0.02% (v/v) Tween 20 as a surfactant. After 48 hours, treated leaves were immersed for 24 hours in 40 mL of conidia solution which contained 2×10^4^ conidia/mL before the conidia solution was removed. The progression of the infection was evaluated by the quantification of *C. graminicola* genomic DNA by quantitative PCR. Maize leaf discs sampled at 4 dpi were homogenized using metal beads in a Retsch mixer mill and fungal genomic DNA was extracted with NucleoSpin® Plant II Kit (Macherey-Nagel, Dueren, Germany) as explained before. *C. graminicola* genomic DNA was quantified by quantitative PCR using Biozym Blue S’Green qPCR Kit (Biozym Scientific, Germany) in a CFX RT-PCR detection system (Bio-Rad, USA) and specific primers for the *histone 3* (Cg*H3*) gene (Supplementary Table S1). The relative quantity of *C. graminicola* genomic DNA was normalized to leaf area (cm^2^).

Wheat plants were grown for 2 weeks and whole plants were sprayed with 0.80 mL per plant of water (mock) or 500 µM B2G3, both solutions supplemented with 0.1% (v/v) Tween 20. After 24 hours, plants were sprayed with 12.5 mL of *Z. tritici* strain ST99CH_3D7 spore suspension which contained 1×10^7^ spores/mL and 0.1% (v/v) Tween 20. Pots were enclosed with a plastic bag for 72 hours to ensure high humidity. The percentage of leaf area covered by lesions and the number of pycnidia per cm^2^ of leaf were assessed 12 dpi. Sixteen second leaves per treatment were mounted on paper sheets, scanned using a CanoScan LiDE 400 scanner, and analysed with ImageJ (Schneider et al., 2012) and an automated image analysis method (Stewart et al. 2016).

### Data analysis and software

For dose-response analyses, total relative luminescence units (RLU) were obtained by calculating the integral under the kinetic curve in each case. The estimated effective dose (ED) was calculated using the total RLUs and an “R” script using the package drc v3.0 as described by Ritz et al. (2015). R software (v4.2.2, R Development Core Team, 2008; https://www.r-project.org/) was used for calculating the ED and drawing the figure.

For statistical analysis, Student’s t-test was used to determine whether or not a set of data was significantly different compared to the mock, n.s. means non-significant differences (p>0.05) and asterisks indicate significant differences (*p ≤ 0.05, **p ≤ 0.01, ***p ≤ 0.001). The results were analysed using SPSS statistical software (version 29.0.2.0).

## Results

### β-1,2-glucan oligosaccharides trigger H_2_O_2_ production in Arabidopsis plants in a dose-dependent manner

One of the first PTI responses in plants is the quick production of ROS (mainly H_2_O_2_) upon the recognition of a molecular pattern. Therefore, the luminol-based assay to quantify ROS production after treatment with peptides or oligosaccharides has been extensively used to identify novel plant defence elicitors. The treatment of Arabidopsis seedlings with 100 µM α-1,3-glucan trisaccharide (A3G3) had no effect on ROS production, which, to our knowledge, had not been demonstrated before. However, β-1,2-glucan oligosaccharide treatments (100 µM B2G3-trisaccharide and B2G6-hexasaccharide; Supplementary Fig. S1) triggered the production of ROS following a kinetics similar to that of previously described plant defence elicitors such as MLG43 and Chitohexaose (Chi6) (Fig. 1A). In fact, the quantification of the total amount of RLUs showed a significant increase in ROS production in Chi6, MLG43, B2G3 and B2G6 oligosaccharides in comparison with A3G3- and water-treated plants (mock) (Fig. 1B). In contrast, treatment with a β-1,2-glucan polysaccharide led to a slight, but non-significant, increase in ROS production at the concentrations tested (0.1 to 0.5 mg/mL) (Fig. 1A and B and Supplementary Fig. S2).

**Fig. 1.**
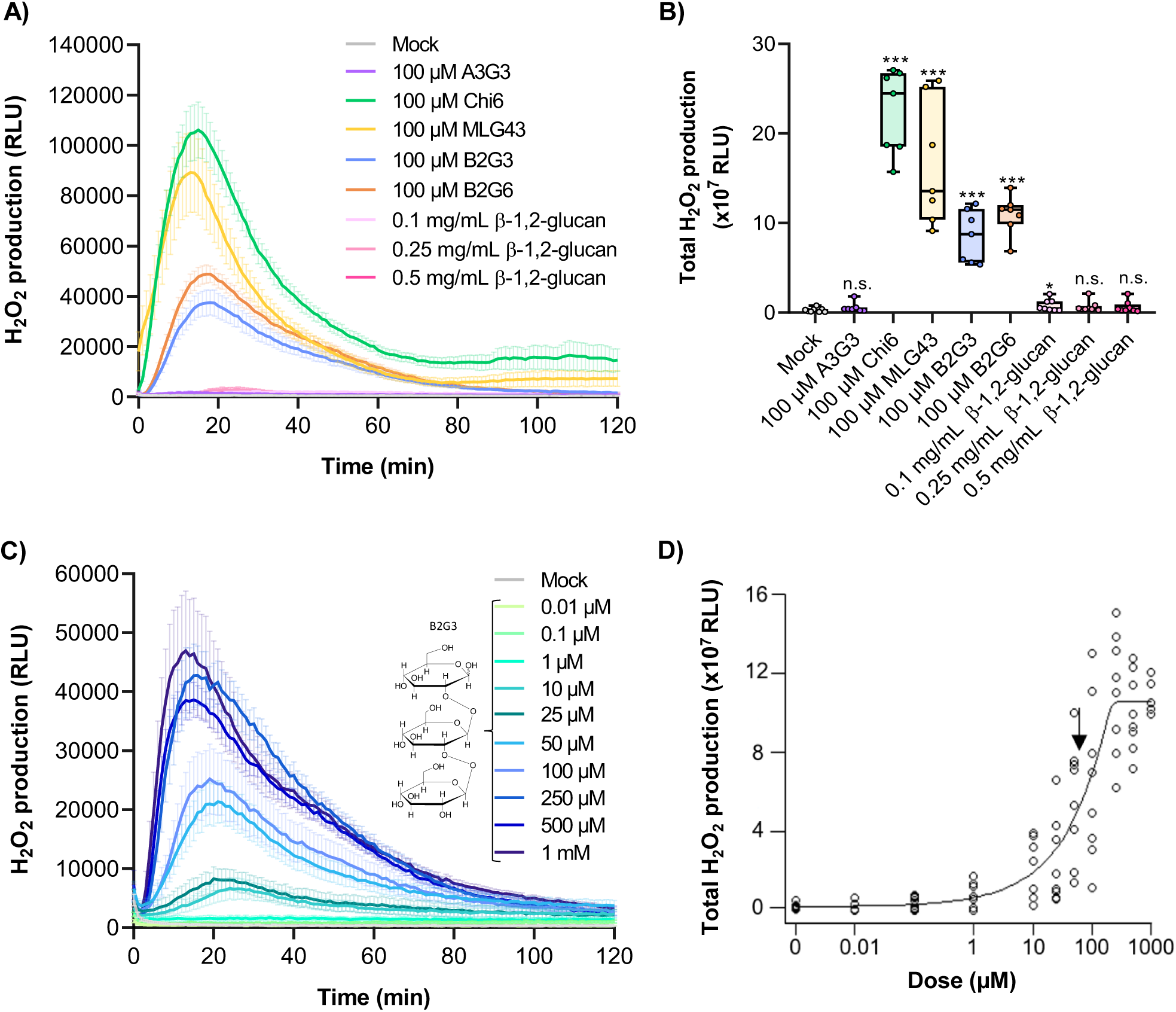
β-1,2-glucan (B2G3 and B2G6) oligosaccharides trigger reactive oxygen species (ROS) production in Arabidopsis seedlings. (A) ROS production in Arabidopsis (Col-0) seedling using luminol reaction measured as relative luminescence units (RLU) over time using β-1,2-glucan oligosaccharides (trimer, B2G3 and hexamer, B2G6 at a concentration of 100 µM), β-1,2-glucan polysaccharide (0.1 to 0.5 mg/mL), Hexaacetyl-Chitohexaose (Chi6, 100 µM), β-D-cellobiosyl-(1,3)-β-D-glucose (MLG43, 100 µM), α-1,3-glucan trisaccharide (A3G3, 100 µM) and water (mock), which was the negative control. (B) Total production of ROS measured as total RLUs over 120 minutes after the treatment with oligosaccharides. Statistically non-significant (n.s.) and significant differences according to Student’s t-test (*p ≤ 0.05, ***p ≤ 0.001) compared to the mock are shown. (C) ROS production using different concentrations of B2G3 (from 0.01 µM to 1 mM). Data represent mean ± standard error (n=8) from one experiment out of three performed that produced similar results. (D) Dose-response curve in Arabidopsis seedlings. Total response measured as total RLUs along 120 minutes after elicitation with different concentrations of B2G3 (from 0.01 µM to 1 mM). The arrow indicates the estimated effective dose (EED; 50% of total signal), 64.83 µM in this case.

To characterize the kinetics of β-1,2-glucan oligosaccharide perception, we treated Arabidopsis seedlings with different concentrations of B2G3 ranging from 0.01 µM to 1 mM. Results indicated that ROS production is dependent on B2G3 concentration, and this oligosaccharide is effective at the micromolar range (from 1 µM to 250 µM B2G3) (Fig. 1C and D). The calculation of the estimated effective dose (EED, 50% of total signal) was 64.83 µM, which is similar to EED values for other glycan-based immune elicitors such as MLG43, Chi6 and β-1,3-glucan hexasaccharide (Mélida et al., 2018; Rebaque et al., 2021) (Fig. 1D).

### B2G3 and B2G6 activate other PTI hallmarks

To confirm that β-1,2-glucan oligosaccharides effectively trigger PTI responses, we evaluated the phosphorylation of MAPKs (MAPK3, MAPK6 and MAPK4/11) by western-blot using an anti-p44/42 antibody which specifically recognizes the phosphorylated forms of these MAPKs. As previously described, MLG43 (100 µM), used as positive control, induced MAPK3 and MAPK6 phosphorylation after 5 minutes of treatment, but MAPK4/11 phosphorylation was not observed (Fig. 2A and B; Rebaque et al., 2021). Interestingly, 100 µM B2G3 and B2G6 showed a peak of MAPK phosphorylation at 15 minutes after treatment (Fig. 2A and B). B2G3 treatment showed the phosphorylation of all MAPKs (MAPK3, MAPK6 and MAPK4/11), whereas B2G6 only induced the phosphorylation of MAPK6, but not of MAPK3 and MAPK4/11.

**Fig. 2.**
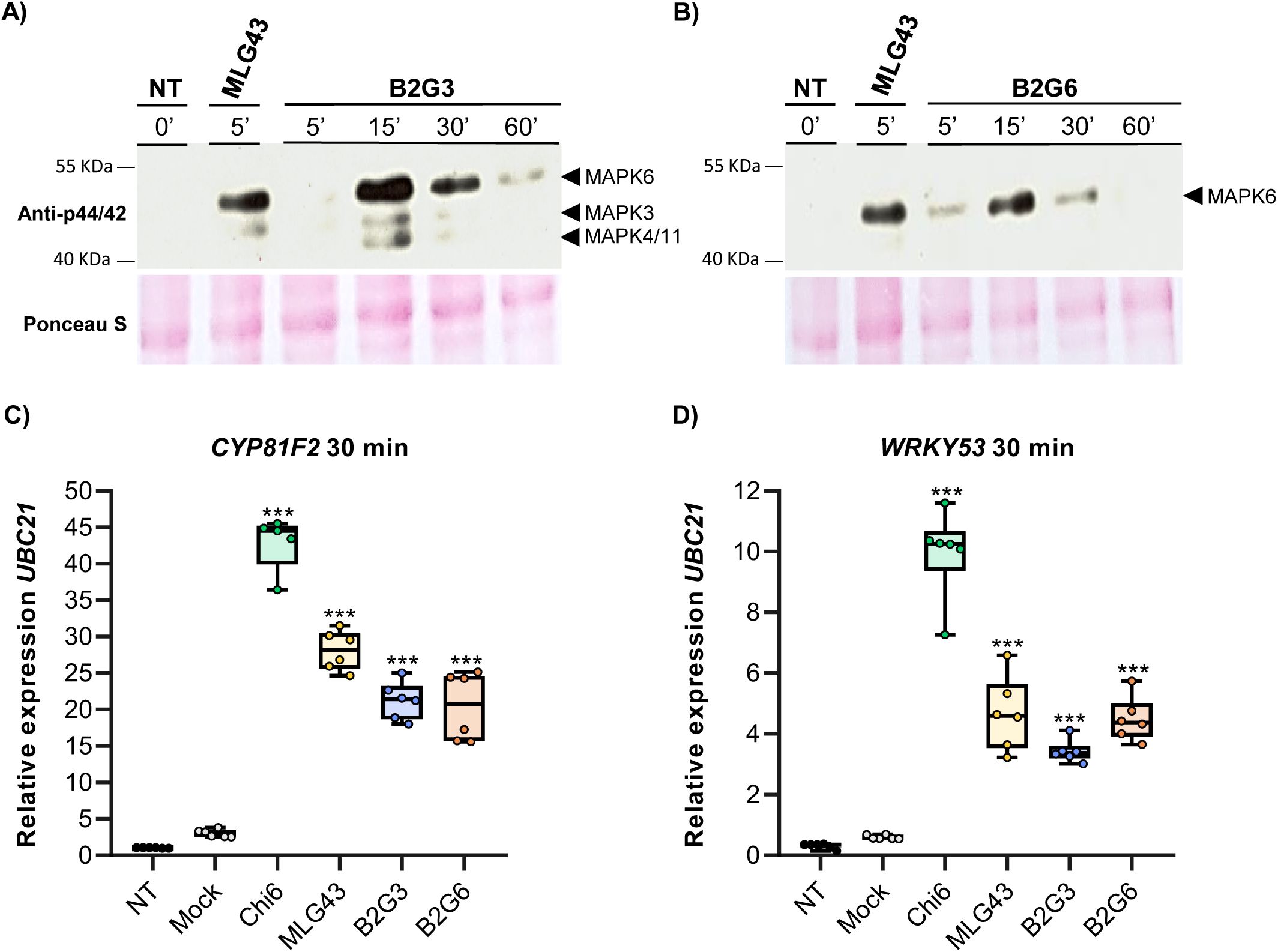
Pattern-triggered immunity hallmark activation by B2G3 and B2G6 in Arabidopsis. Mitogen-activated protein kinases (MAPK) phosphorylation in seedlings determined by Western Blot using Phospho-p44/42 MAPK antibody for phosphorylated MAPK moieties at different time points (5, 15, 30 and 60 minutes) after treatment with (A) B2G3 (100 µM) and (B) B2G6 (100 µM). NT means non-treated sample, which was the negative control, and MLG43 (100 µM), was used as positive control. Arrowheads indicate the position of phosphorylated MPK6 (top), MPK3 (middle) and MPK4/11 (bottom). Ponceau S red-stained membranes show equal protein loading. Quantitative RT-PCR analysis of the relative expression levels of immune-related genes (C) *CYP81F2* and (D) *WRKY53* normalized to the expression of *UBC21* gene at 30 minutes in treated and non-treated Arabidopsis seedlings (n = 6). NT and distilled water (mock) were used as negative controls and Chi6 and MLG43 were used as positive controls. All treatments were performed at 100 µM. Data are presented as box plots, with the centre line showing the median, the box limits showing the 25^th^ and 75^th^ percentiles, and the whiskers showing the full range of data. All the data displayed belong to one of the three independent experiments carried out, which gave similar results. Statistically significant differences according to Student’s t-test (***p ≤ 0.001) compared to the mock are shown.

In addition, we also evaluated the expression of two PTI reporter genes, *CYP81F2* and *WRKY53,* after 30 minutes of treatment with oligosaccharides. *CYP81F2* (*CYTOCHROME P450, FAMILY 81*) encodes a cytochrome P450 monooxygenase involved in the biosynthesis of indole glucosinolates, while *WRKY53* (*WRKY DNA-BINDING PROTEIN 53*) is involved in basal resistance against *P. syringae* (Murray et al., 2007; Pfalz et al., 2009). As expected, Chi6 and MLG43 treatments strongly induced the expression of both PTI reporter genes (Fig. 2C and D). Similarly, β-1,2-glucan oligosaccharides, B2G3 and B2G6, also led to a significant increase in the expression of both PTI reporter genes (Fig. 2C and D). Taken together, both β-1,2-glucan oligosaccharides triggered all PTI responses evaluated, but B2G3 caused the phosphorylation of all MAPKs. For this reason, this oligosaccharide was selected for further experiments.

### B2G3 treatment reduces Arabidopsis disease symptoms caused by different pathogens

It has been extensively described that linear or cyclic β-1,2-glucan polymers (with a degree of polymerization from 6 to 40 glucoses) are deposited in the periplasmic space of certain groups of Gram-negative plant-colonizing bacteria as osmoregulated periplasmic glucans (OPGs) and play multiple roles in the bacterial lifestyle (Bohin, 2000; Wanke et al., 2021). Considering this and that β-1,2-glucan oligosaccharides are capable of triggering PTI responses, we wondered if B2G3 pretreatment would protect plants after the inoculation of pathogenic bacteria such as *Pseudomonas syringae* pv. *tomato* (DC3000). Bacterial inoculation produced the appearance of chlorotic lesions on rosette leaves of mock-pretreated Arabidopsis plants at 3, 5 and 7 dpi, while the pretreatment with B2G3 strongly reduced those symptoms (Fig. 3A). These results are in accordance with the significant reduction of DC3000 growth at 3 dpi in B2G3 pretreated plants in comparison with mock-pretreated plants (Fig. 3B). In conclusion, B2G3 is effective in the activation of PTI responses that protect against *P. syringae* in Arabidopsis.

**Fig. 3.**
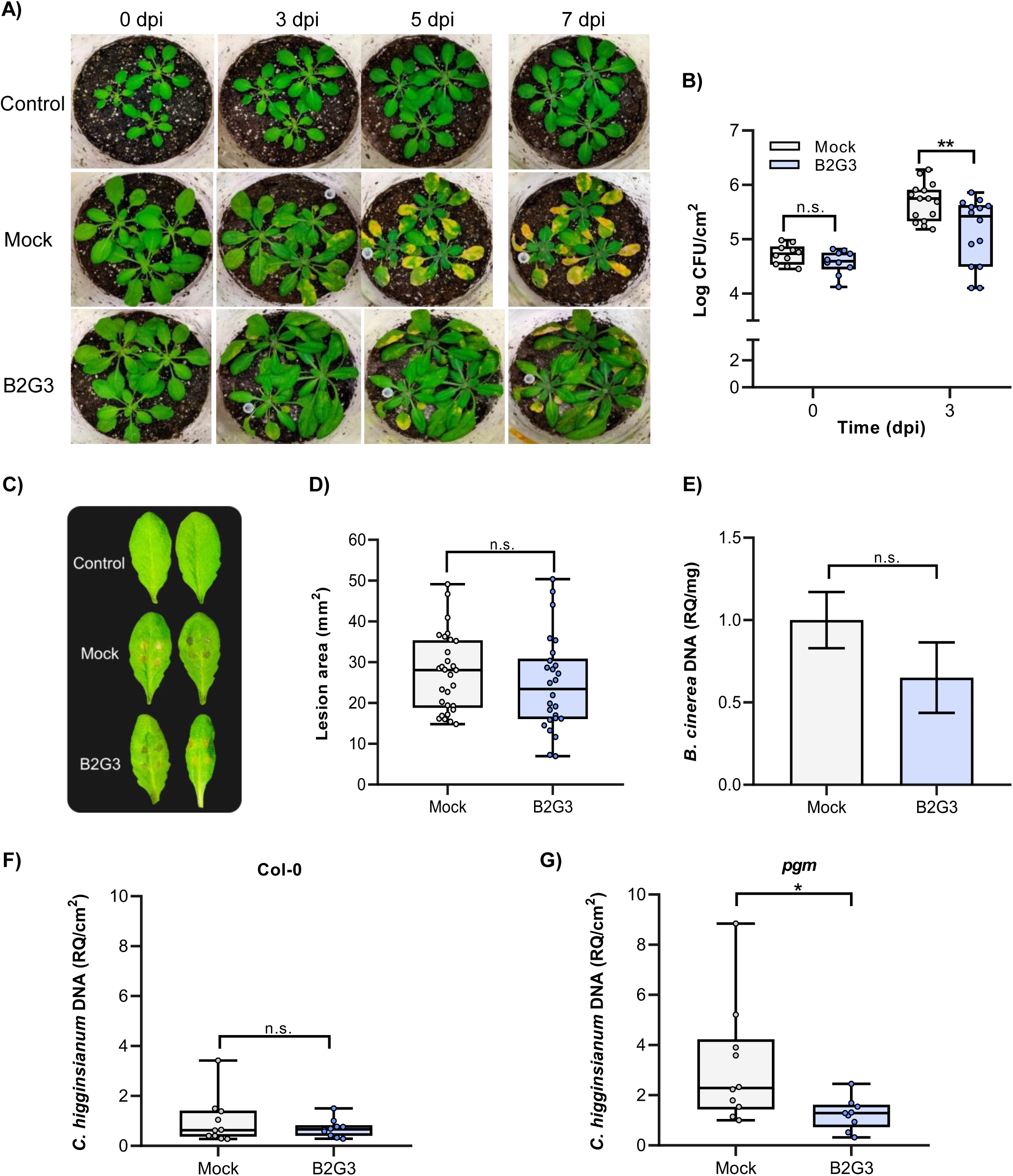
Treatment of Arabidopsis plants with B2G3 enhance protection against phytopathogenic bacteria. (A) Plants of Arabidopsis (Col-0) pretreated with distilled water (mock) or B2G3 (500 µM and 0.5 mL per plant) and inoculated with *Pseudomonas syringae* pv. *tomato* (DC3000). Control refers to non-inoculated plants. Pictures were taken from 0 to 7 days post-infection (dpi) and representative images are displayed. (B) Colony forming units (CFU) of DC3000 per leaf area (cm^2^) at 0 and 3 dpi (n=10-13). (C) Disease symptoms produced by *Botrytis cinerea* at 2 dpi in Arabidopsis leaves. These plants were non-pretreated or pretreated with distilled water (mock) or B2G3 (500 µM and 0.5 mL per plant). (D) Lesion area (mm^2^) caused by *B. cinerea* infection at 2 dpi in Arabidopsis plants (n=26-29, from 15 different plants). (E) Quantification of *B. cinerea* genomic DNA in inoculated Arabidopsis leaves at 1 dpi by quantitative PCR using specific primers for the *B. cinerea* β*-Tubulin* (Bc*Tub*) gene. The genomic DNA quantification is expressed as relative quantity (RQ) to Arabidopsis leaves weight (mg) (n=3). Quantification of *Colletotrichum higginsianum* genomic DNA in Arabidopsis (F) Col-0 and (G) the *C. higginsianum*-hypersensusceptible mutant *pgm* after treatment with distilled water (mock) and B2G3 (1 mM and 500 µL per plant). Genomic DNA was quantified at 3.5 dpi by quantitative PCR using specific primers for *C. higginsianum TrpC* amplification and data is represented as RQ of genomic DNA per leaf area (cm^2^) (n=9-10). Statistically non-significant (n.s.) and significant differences according to Student’s t-test (*p ≤ 0.05, **p ≤ 0.01) compared to the mock are shown.

The occurrence of β-1,2-glucans has been previously described in fungal and oomycete cell walls (Mitchell and Sabar, 1966; Ruiz-Herrera and Ortiz-Castellanos, 2019). Therefore, we decided to test if β-1,2-glucan oligosaccharides protect Arabidopsis plants against fungal diseases. To achieve this, we treated Arabidopsis plants with B2G3 prior to inoculation of two fungal pathogens, the necrotrophic *Botrytis cinerea* and the hemibiotrophic *Colletotrichum higginsianum*. B2G3 treatment did not reduce the necrotic lesion area of *B. cinerea* at 2 dpi after droplet inoculation on Arabidopsis leaves (Fig. 3C and D). Quantification of the genomic DNA of *B. cinerea* in infected leaves after 24 hours of infection also showed that the pretreatment with B2G3 had no effect on the growth of this fungus (Fig. 3E). For *C. higginsianum* infection experiments, we pretreated Arabidopsis plants with B2G3 and performed a second B2G3 treatment at the time of infection. Nevertheless, the amount of the *C. higginsianum* genomic DNA in infected wild type leaves was similar to mock-treated plants at 3.5 dpi (Fig. 3F). To investigate if B2G3 treatment can protect hypersusceptible Arabidopsis plants, we performed the same treatments on the starch-deficient *pgm* mutants, which are hypersusceptible to this fungal infection due to reduced carbohydrate availability and reduced penetration resistance (Engelsdorf et al., 2013, 2017). Treatment of *pgm* plants with B2G3 caused a significant reduction in fungal genomic DNA at 3.5 dpi, indicating that plants with impaired basal resistance can be protected by B2G3 application (Fig. 3F). These results would classify B2G3 as a new elicitor able to trigger PTI responses in Arabidopsis, which led to protection against certain pathogens.

### B2G3 trigger PTI immune responses and confers defence against pathogens in monocot plants

We further evaluated the potential of B2G3 in triggering PTI in plants other than Arabidopsis. In order to evaluate if β-1,2-glucans trigger an immune response in maize plants, we first determined the production of H_2_O_2_ in maize leaf discs after treatment with these glycans. Treatment with B2G3 and B2G6 (100 µM) significantly triggered ROS production in maize discs in a similar way as Chi6 (100 µM) (Fig. 4A). Total RLU quantification showed a significant increase in ROS production after Chi6, B2G3 and B2G6 treatment in comparison with mock-treated plants (Fig. 4B). As observed in Arabidopsis seedlings, A3G3 treatment did not trigger ROS production (Fig. 4A and B). To assess whether the PTI defence responses triggered by B2G3 protected maize plants against diseases, we pretreated maize plants with MLG43 and B2G3 prior to inoculation with the hemibiotrophic fungal pathogen *Colletotrichum graminicola*. Pretreatment of maize plants with MLG43 or B2G3 reduced fungal growth, as evidenced by a significant reduction in the amount of *C. graminicola* genomic DNA at 4 dpi and a reduction of fungal development during the pathogenesis (Fig. 4C, Supplementary Fig. S3), demonstrating that B2G3 protects maize plants against this fungal pathogen.

**Fig. 4.**
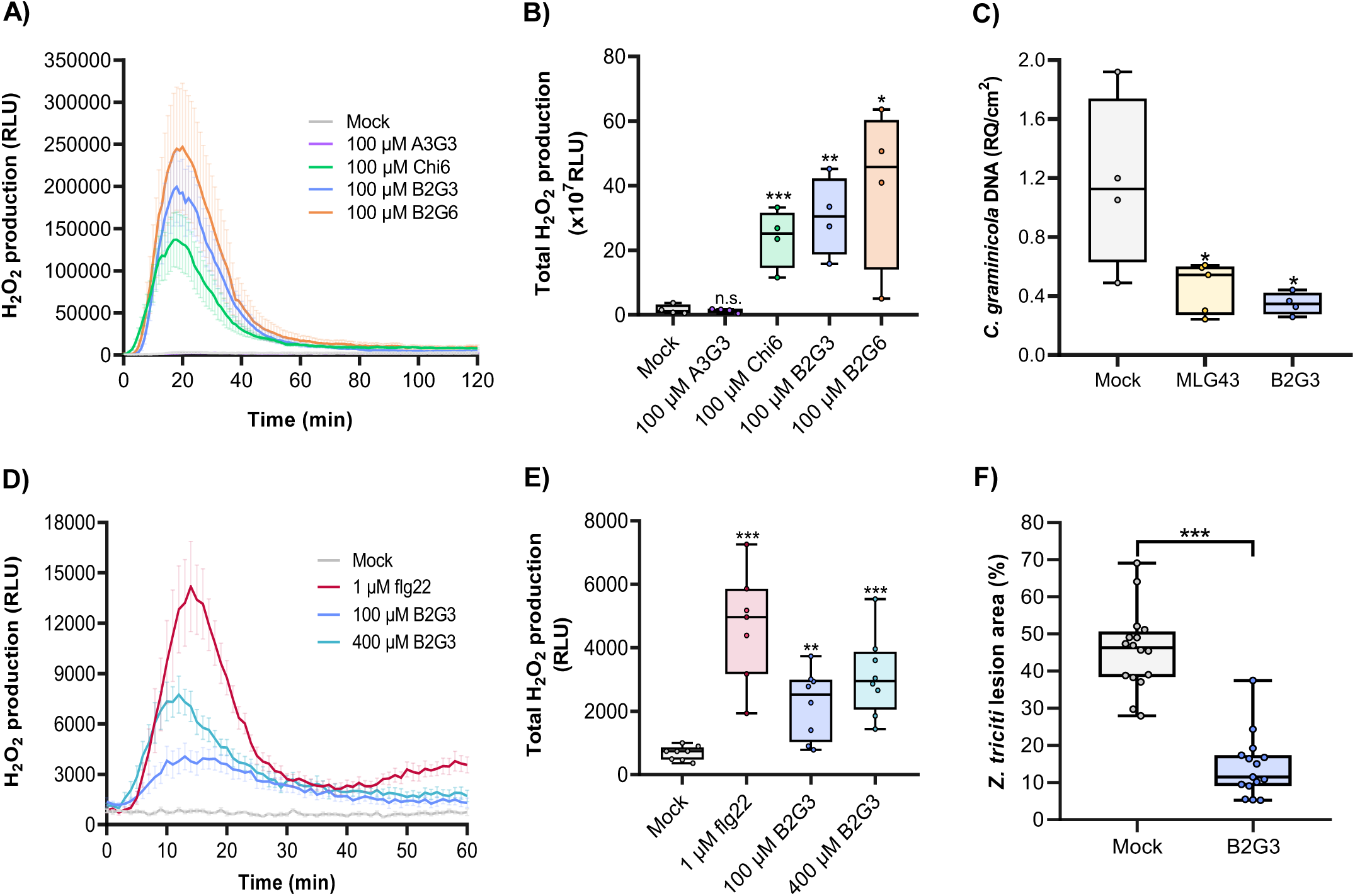
B2G3 induces an immune response in maize and wheat and protects against fungal diseases. (A) Reactive oxygen species (ROS) production in maize plants using luminol reaction measured as relative luminescence units (RLU) over time after treatment with A3G3, Chi6, B2G3 and B2G6 (all of them at 100 µM). Distilled water (mock) was used as negative control. Data represent mean ± standard error (n=8). (B) Total ROS production in maize plants measured as total RLUs over 120 minutes after treatment with the oligosaccharides. (C) Quantification of *Colletotrichum graminicola* genomic DNA per maize leaf area (cm^2^). Genomic DNA was quantified at 4 dpi by quantitative PCR using *C. graminicola* specific primers for histone 3 (Cg*H3*) (n=4-5). The experiment was performed in plants treated with distilled water (mock), MLG43 (1 mM and 0.5 mL per plant) (positive control) and B2G3 (1 mM and 0.5 mL per plant). (D) ROS production in wheat plants using luminol reaction measured as RLU over time after treatment with flg22 (1 µM) and two concentrations of B2G3 (100 and 400 µM). Distilled water (mock) was used as negative control. Data represent mean ± standard error (n=8). (E) Total ROS production in wheat plants measured as total RLUs along 60 minutes after treatment with flg22 (1µM) and B2G3 (100 and 400 µM). (F) Lesion area (%) caused by *Zymoseptoria tritici* infection in wheat plants after treatment with distilled water (mock) and B2G3 (500 µM and 0.78 mL per plant) (n=16) at 12 dpi. Statistically non-significant (n.s.) and significant differences according to Student’s t-test (*p ≤ 0.05, **p ≤ 0.01, ***p ≤ 0.001) compared to the mock are indicated.

We also explored the effect of β-1,2-glucan trisaccharide treatment on wheat plants. As previously observed for Arabidopsis and maize, B2G3 induced a significant increase in H_2_O_2_ production, which confirmed that B2G3 was perceived by wheat PRRs (Fig. 4D and E). We further investigated whether B2G3 protects wheat against pathogen diseases by infecting B2G3-pretreated wheat plants with *Zymoseptoria tritici*, a latent fungal necrotroph (Sánchez-Vallet et al., 2015). B2G3 pretreatment significantly reduced symptom development produced by *Z. tritici*, as shown by the restricted lesion area and the significant reduction in pycnidia density in B2G3-pre-treated plants (Fig. 4F and Supplementary Fig. S4). Thus, our results demonstrated that B2G3 triggered the immune system in crops such as maize and wheat and protect them against fungal diseases.

## Discussion

Although glucans were the first group of plant elicitors proposed as such in the 1970s thanks to pioneering works by several groups (Ayers et al., 1976a,b; Anderson, 1978), it is only recently, with the resurgence of research into glycan-triggered plant immunity, that a better understanding of their ability to induce such defensive responses has begun to emerge (Molina et al., 2024). In fact, there have been several groups in the current century that have made significant advances in the field. Klarzynski et al. (2000) discovered that linear unsubstituted β-1,3-glucan oligosaccharides extracted from algae cell walls trigger immune responses in tobacco (*Nicotiana tabacum*), that protected from soft rot pathogen *Erwinia carotovora*. Similarly, cellodextrins, water-soluble β-1,4-glucan oligosaccharides derived from cellulose, induced immune responses in gravepine (*Vitis vinifera*) cells and reduced *B. cinerea* infection symptoms when applied to leaves prior fungal inoculation (Aziz et al., 2007). Despite these advances, it was even proposed that the slow progress in the study of glucan-triggered immunity was due to the insensitivity of the Arabidopsis ecotype Col-0 to glucans (Fesel and Zuccaro, 2016). However, numerous studies published over the last decade have shown that Arabidopsis is a good system to study glucan-triggered immunity (Souza et al., 2017; Claverie et al., 2018; Johnson et al., 2018; Mélida et al., 2018; Locci et al., 2019; Wanke et al., 2020; Rebaque et al., 2021, 2023; Zarattini et al., 2021; Tseng et al., 2022; Martín-Dacal et al., 2023). The studies of two research groups confirmed the data published by Aziz et al., (2007) a decade earlier on the ability of β-1,4-glucans to induce PTI responses in plants (Souza et al., 2017; Johnson et al., 2018). In fact, this is the only group of β-glucans for which a *bona-fide* interaction with a PRR receptor has been demonstrated to date (Martín-Dacal et al., 2023). Shortly afterwards it was also confirmed that β-1,3-glucans induced defensive responses in Arabidopsis and barley (*Hordeum vulgare*), and it was proposed that these responses were dependent on the CERK1 coreceptor in the case of the hexasaccharide in Arabidopsis (Mélida et al., 2018; Wanke et al., 2020). Subsequently, a group of glucans containing both β-1,3 and β-1,4 linkages, also known as mixed-linked glucans (Burton and Fincher, 2009), were proposed as a very promising group of plant immunostimulants, inducing potent defence responses at low doses (Barghahn et al., 2021; Rebaque et al., 2021; Yang et al., 2021). Furthermore, β-1,6-glucans have also recently been shown to induce phosphorylation of MAPKs in Arabidopsis, although their protective capacity against diseases has not been investigated yet (Chaube et al., 2022; Fernández-Calvo et al., 2024). Thus, glucans have positioned themselves as a core group regulating glycan-triggered immunity in plants. The fact that glucans-based elicitors have been evolutionarily selected as such makes sense, given that these carbohydrates are present in the extracellular matrices of all microorganisms that interact with plants (Mélida et al., 2013; Bontemps-Gallo and Lacroix, 2015; Ruiz-Herrera and Ortiz-Castellanos, 2019; Yugueros et al., 2024). However, there are two groups of glucans that have hardly been studied in the field of plant immunity, β-1,2- and α-1,3-glucans, which are also present in these extracellular matrices (Bohin, 2000; Bontemps-Gallo and Lacroix, 2015; Ruiz-Herrera and Ortiz-Castellanos, 2019).

Linear α-1,3-glucans are one of the major components of fungal cell walls, however, their occurrence is specific of Dikarya subkingdom, being absent in lower fungal phyla (Yoshimi et al., 2017; Ruiz-Herrera and Ortiz-Castellanos, 2019). Although the deposition of this polysaccharide close to cell membranes has been described, it is frequently exposed in the most extracellular layer of fungal walls, acting as an aggregation factor during hyphae and conidia development (Yoshimi et al., 2017; Sugawara et al., 2003). As an exposed glucan on mammal pathogenic fungal cell walls, research has demonstrated their role as virulence factors acting as immunosuppressors by blocking the recognition of immunogenic β-1,3-glucans by dectin-1 receptor (Beauvais et al., 2013; Rappleye et al., 2004, 2007). Indeed, there are no reports describing their role as elicitors of animal defence responses (Yoshimi et al., 2017; Ruiz-Herrera and Ortiz-Castellanos, 2019). Although there is low information about the role of α-1,3-glucans in plant defence responses, it has been proposed that these glucans cover the surface of the cell wall of the rice fungal pathogen *Magnaporthe oryzae* and are essential for the infection by protecting the hyphae from host recognition (Fujikawa et al., 2009, 2012). However, the ability of α-1,3-glucan oligosaccharides to trigger plant immune responses has not been reported yet. Here, we demonstrate that, at least, the α-1,3-glucan trisaccharide (A3G3) is not able to trigger the production of ROS after treatment of Arabidopsis seedlings and maize plants, suggesting that this glucan derivative did not trigger plant defence responses.

The occurrence of β-1,2-glucans has been proposed in the cell walls of the green microalgae *Chlorella pyrenoidosa* (Suárez et al., 2008) and in plant-interacting organisms such as fungi and oomycetes (Mitchel and Sabar, 1966; Ruiz-Herrera and Ortiz-Castellanos, 2019). However, glucans with this particular β-1,2-linkage are one of the main components of OPGs that are deposited in the periplasmic space of Gram-negative symbiotic and pathogenic proteobacteria (Bohin, 2000; Bontemps-Gallo & Lacroix, 2015; Rigano et al., 2007). They are composed of linear or cyclic β-1,2-glucans with different degree of polymerization and diverse β- and α-linked glucose substitutions depending on the group of proteobacteria considered (Bohin, 2000; Bontemps-Gallo and Lacroix, 2015). These polymers play multiple roles during bacteria development such as motility, cell division, sensitivity to antibiotics and biofilm formation, and they are essential for maintaining full virulence in animal and plant pathogenic bacteria by indirectly controlling virulence-gene expression (Bohin, 2000; Bontemps-Gallo et al., 2013; Bontemps-Gallo and Lacroix, 2015). Contrarily to A3G3, β-1,2-glucan trisaccharide (B2G3) and hexasaccharide (B2G6) successfully triggered several PTI hallmarks in Arabidopsis at low concentrations, such as ROS production, the phosphorylation of defence-related MAPKs (MAPK3, MAPK6 and MAPK4/11) and the overexpression of PTI-related genes. The effect on PTI hallmarks of B2G3 and B2G6 is quite similar to glycan-based defence elicitors MLG43 and Chi6 (Mélida et al., 2018; Rebaque et al., 2021; Shi et al., 2019), confirming their role as plant immunostimulants. Moreover, we determined the dose-dependence of B2G3 in Arabidopsis seedlings in order to trigger PTI responses and showed that this oligosaccharide is active at micromolar range, which is a similar dose as that of other glucan oligosaccharides such as xyloglucan and β-1,3-, β-1,4- and β-1,3/1,4-linked-glucans (Klarzynski et al., 2000; Aziz et al., 2007; Claverie et al., 2018; Mélida et al., 2018; Rebaque et al., 2021). In addition, one of the most interesting results that we found is that this PTI induction resulted in increased resistance to *P. syringae*. However, a different scenario was found for fungal pathogens, as B2G3 was not successful in the protection of Arabidopsis wild type plants against the fungi tested. In this case, we challenged Arabidopsis B2G3-pretreated plants with *B. cinerea* and *C. higginsianum*, which are characterised by necrotrophic and hemibiotrophic infection styles against Arabidopsis under laboratory conditions (Engelsdorf et al., 2013; Zarattini et al., 2021). In this context, it has been shown that fungi have developed mechanisms to hide elicitor molecules derived from their cell wall by combining them with lectins or other types of effectors (Wawra et al., 2016, 2019; Irieda et al., 2019; Sánchez-Vallet et al., 2020). Nevertheless, we could observe significant protection by this trisaccharide when we challenged hypersusceptible plants with *C. higginsianum*. These results open the door to investigate the molecular mechanisms controlling β-1,2-glucan-mediated fungal disease tolerance in Arabidopsis.

As they represent important components of symbiotic and pathogenic bacteria, β-1,2-glucans are good candidates for immunomodulation of defence responses in host plants, but to date there is only one report that demonstrate the ability of cyclic β-1,2-glucans from *Xanthomonas campestris* pv *campestris* to suppress systemically plant immune responses in Arabidopsis and *Nicotiana benthamiana* (Rigano et al., 2007). At first sight, these results seemed contradictory to the results reported here, but it has been demonstrated that the immunomodulatory effect of glycan oligomers is dependent on their structure, being especially important the DP and the presence/absence of different substitutions Klarzynski et al., 2000; Mélida et al., 2018; Voxeur et al., 2019; Chandrasekar et al., 2022). This is the case for non-branched β-1,3-glucans oligomers whose DP determine the ability of triggering PTI responses and the substitution with β-1,6-linked glucoses depleted their immunostimulatory activity (Klarzynski et al., 2000; Mélida et al., 2018). However, a recent work highlights that a β-1,3/1,6-glucan decasaccharide naturally obtained after the action of a barley β-1,3-endoglucanase, is not able to activate immune defences, but scavenge ROS, enhancing pathogen colonization (Chandrasekar et al., 2022). Another interesting mechanism of activation/inactivation of glucan-eliciting capacity is that mediated by the berberine bridge enzyme-like oxidases of β-1,4- and β-1,3/1,4-linked β-glucans (Locci et al., 2019; Costantini et al., 2024). These enzymes, through simple oxidation, deactivate the immunostimulatory capacity of these glucans, which has been proposed as a homeostatic control mechanism to prevent the hyperaccumulation of elicitors. As the above examples illustrate, there is fine control in the mechanisms of glycan-mediated immunostimulation by both host and pathogen. It is therefore possible that β-1,2-glucans have the dual ability to suppress or stimulate plant defences, depending on the form in which they are presented to the plants. Therefore, enzymatic cleavage of cyclic glucans of *X. campestris* (15 β-1,2-linkages and one α-1,6-linkage) could lead to immune active β-1,2-glucan oligomers, a hypothesis that is reinforced by the description of bacterial and fungal β-1,2-endoglucanases that generate B2G3 and B2G6 as products (Abe et al., 2017; Tanaka et al., 2019).

Furthermore, our results obtained in Arabidopsis are reinforced by those generated with two monocotyledonous species, maize and wheat. We demonstrated that β-1,2-glucan oligosaccharides treatment caused ROS accumulation in maize and wheat similar to glycan oligosaccharides derived from chitin and mixed-linked glucans or the peptide MAMP flg22. These results indicate that both plants have the machinery of perception of these β-1,2-glucan oligomers. Interestingly, the activation of plant PTI responses led to enhanced disease resistance to *C. graminicola* (maize) and *Z. tritici* (wheat). In fact, to date there are hardly any studies on glycan-induced immunity in this group of plant species. On the other hand, taking into account that the few studies that exist indicate that there are certain differences between the mechanisms of glycan perception by PRRs between monocots and dicots (Liu et al., 2016; Wanke et al., 2020; Yang et al., 2021), it will be interesting to continue to deepen our knowledge of the mechanisms involved in the perception of this new group of elicitors described in this work.

In summary, our results demonstrate that short β-1,2-glucan oligosaccharides led to the activation of PTI responses that enhanced disease resistance to bacterial and fungal pathogens in phylogenetically distant plants. Therefore, this work supports the plant immunostimulatory capacity of a previously uncharacterized group of β-glucans and opens the way to future research into the molecular mechanisms controlling β-glucan recognition, as well as the development of plant disease control methods based on β-glucan application.

## Supplementary data

Table S1. Primers (Forward (Fw) and Reverse (Rv)) used for gene expression analyses.

Fig. S1. Structural scheme of the β-1,2-oligosaccharides studied.

Fig. S2. β-1,2-glucan trigger reactive oxygen species (ROS) production in Arabidopsis seedlings.

Fig. S3. Maize leaf pretreatment with MLG43 and B2G3 delays the development of *Colletotrichum graminicola*.

Fig. S4. Treatment of wheat plants with B2G3 enhance protection against *Zymoseptoria tritici*.

## Author contribution

HM: conceptualization; MFR, ALG, AF, KM and CCL: investigation; ALG, ASV, TE and HM: supervision; MFR, ALG and HM: writing – original draft preparation; MFR, ALG, AF, KM, CCL, ASV and HM: writing - review & editing.

## Data availability

Data will be made available on request.

## Supplementary data

**Table S1.**
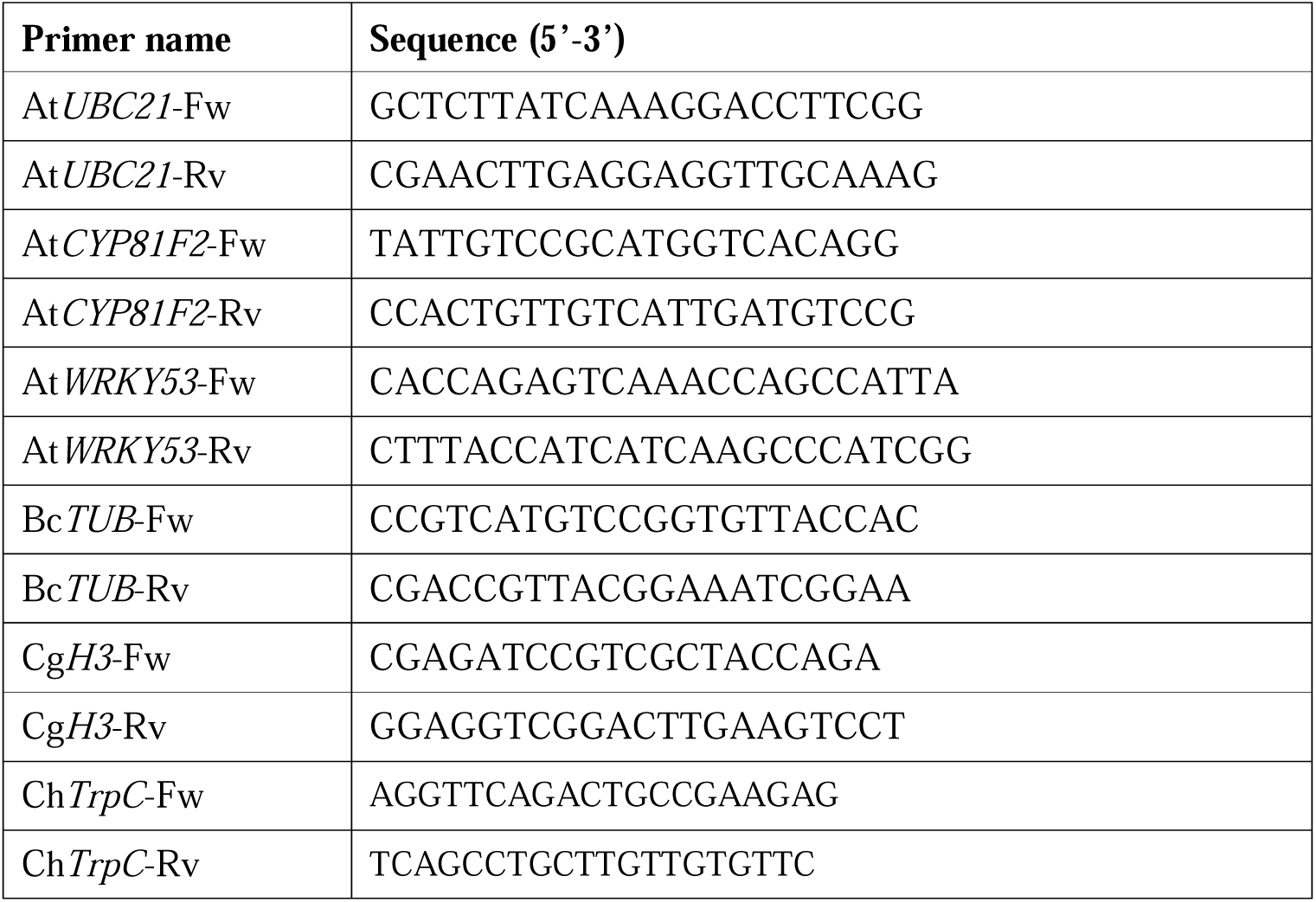
Primers (Forward (Fw) and Reverse (Rv)) used for gene expression analyses.

**Fig. S1.** Structural scheme of the β-1,2-oligosaccharides studied. (A) B2G3 and (B) B2G6.

**Fig. S2.** β-1,2-glucan trigger reactive oxygen species (ROS) production in Arabidopsis seedlings. ROS production in Arabidopsis (Col-0) seedling using luminol reaction measured as relative luminescence units (RLU) over time using a mixture of β-1,2-glucan oligosaccharides and water (mock).

**Fig. S3.** Maize leaf pretreatment with MLG43 and B2G3 delays the development of *Colletotrichum graminicola*. (A) Representation of the main structures of *C. graminicola* during pathogenesis of maize leaves at 2.5 dpi. Fungal structures were stained with acid fuchsin and photographed under the microscope. S: ungerminated spores; A: formation of melanized appressoria; PH: development of primary hyphae; SH: development of secondary hyphae. Bar: 20 mm. (B) Proportion of fungal structures on maize leaves after oligosaccharide pretreatment. The quantification was performed by counting more than 200 fungal structures of five biological replicates and indicates to most advanced infection structure per infection event (n=5). Data represent mean ± standard error.

**Fig. S4.** Treatment of wheat plants with B2G3 enhance protection against *Zymoseptoria tritici.* Quantification of *Z. tritici* pycnidia per leaf area (cm^2^) after treatment of wheat plants with distilled water (mock) and B2G3 (500 µM and 0.78 mL per plant) (n=16) at 12 dpi. Statistically significant differences according to Student’s t-test (***p ≤ 0.001) compared to the mock are indicated.

